# A Gold Standard for Transcription Factor Regulatory Interactions in *Escherichia coli* K-12: Architecture of Evidence Types

**DOI:** 10.1101/2023.02.25.530038

**Authors:** Paloma Lara, Socorro Gama-Castro, Heladia Salgado, Claire Rioualen, Víctor H. Tierrafría, Luis J. Muñiz-Rascado, César Bonavides-Martínez, Julio Collado-Vides

## Abstract

Post-genomic implementations have expanded the experimental strategies to identify elements involved in the regulation of transcription initiation. As new methodologies emerge, a natural step is to compare their results with those from established methodologies, such as the classic methods of molecular biology used to characterize transcription factor binding sites, promoters, or transcription units.

In the case of *Escherichia coli* K-12, the best-studied microorganism, for the last 30 years we have continuously gathered such knowledge from original scientific publications, and have organized it in two databases, RegulonDB and EcoCyc. Furthermore, since RegulonDB version 11.0 (1), we offer comprehensive datasets of binding sites from chromatin immunoprecipitation combined with sequencing (ChIP-seq), ChIP combined with exonuclease digestion and next-generation sequencing (ChIP-exo), genomic SELEX screening (gSELEX), and DNA affinity purification sequencing (DAP-seq) HT technologies, as well as additional datasets for transcription start sites, transcription units and RNA sequencing (RNA-seq) expression profiles.

Here, we present for the first time an analysis of the sources of knowledge supporting the collection of transcriptional regulatory interactions (RIs) of *E. coli* K-12. An RI is formed by the transcription factor, its positive or negative effect on a promoter, a gene or transcription unit. We improved the evidence codes so that the specific methods are described, and we classified them into seven independent groups. This is the basis for our updated computation of confidence levels, weak, strong, or confirmed, for the collection of RIs. We compare the confidence levels of the RI collection before and after adding HT evidence illustrating how knowledge will change as more HT data and methods appear in the future. Users can generate subsets filtering out the method they want to benchmark and avoid circularity, or keep for instance only the confirmed interactions.

The comparison of different HT methods with the available datasets indicate that ChIP-seq recovers the highest fraction (>70%) of binding sites present in RegulonDB followed by gSELEX, DAP-seq and ChIP-exo. There is no other genomic database that offers this comprehensive high-quality anatomy of evidence supporting a corpus of transcriptional regulatory interactions.

## 1 Introduction

Genomic sciences have strongly affected the landscape of available experimental strategies to identify, on a genomic scale, a variety of genetic elements, such as transcription factor binding sites (TFBSs) and their subset of transcription factor regulatory sites (TFRSs), i.e., those TFBSs with regulatory evidence for a given transcription factor (TF); transcription start sites (TSSs), transcription termination sites (TTSs), as well as transcription units (TUs), all of these in principle under defined growth conditions. A major concern in our curation planning was how to deal with what we saw as a *tsunami* of data coming from high-throughput (HT) methodologies, and how not to inundate and dilute the decades of previous work reflected in the corpus of knowledge supported by classic molecular biology methods. These methods are well appreciated since, as it is well known, they identify individual elements directly.

As mentioned before, we have been for the last 30 years continuously extracting and gathering in RegulonDB and feeding into EcoCyc knowledge from original scientific publications about regulation of transcription initiation and operon organization in *Escherichia coli* K-12. Although we have for years curated HT data, only recently, since RegulonDB version 11.0, have we the updated collections of publicly available genomic HT datasets of binding sites (from ChIP-seq, ChIP-exo, gSELEX and DAP-seq technologies), of TSSs, TTTs, TUs, and normalized RNA-seq expression profiles (1). In our curation work, we have seen that the publications of these types of approaches frequently compare the obtained results with what is known in RegulonDB (2-17). This motivated us to improve our evidence codes to enhance the use of RegulonDB as the “gold standard”. Certainly, evidence codes used for years both in RegulonDB and EcoCyc were not detailed enough to distinguish different methods. For instance, the terms “binding of purified proteins” or “gene expression analysis” did not specify the method.

RegulonDB and EcoCyc accelerate access to knowledge. An example is their use to quickly find the original publications supporting a specific object (for instance, a promoter, or a regulatory site). However, some objects have different properties that are identified by different methods and which may have been described in different publications. For instance, well-characterized regulatory interactions (RIs) require support of the binding of the TF to a specific site in the genome on the one hand, as well as identifying the function of such a TF site in the activation or repression of the regulated promoter. However, for years, we offered all references for each object together. It is only recently that we started to separate the evidence types and the corresponding references from complex objects.

Briefly, the need to easily distinguish objects based on the approach used (i.e., classic vs HT methods), the fact that RegulonDB sites are used as an index to evaluate the performance of novel methods, and the desire to improve the precision in literature access to specific properties of complex objects such as regulatory interactions (RIs) or promoters, motivated us to update the evidence codes behind the knowledge on the regulation of transcription initiation. The new codes distinguish not only the class of methods but also the specific methodology, distinguishing for instance ChIP-seq from ChIp-exo or gSELEX. We began moving in this direction a few years ago, but it is only in this paper that we report these changes that improved knowledge representation in RegulonDB, enabling subsequent analyses such as those shown below.

Once the new evidence types were defined, we reassessed the way they combine to determine the “confidence level” which, based on the set of evidence types behind an object, assigns it as either weak, strong, or confirmed. We have mapped the RIs with the HT-TFBSs collections and added the corresponding HT binding evidence types to the RIs, improving their confidence level. This was updated in the RegulonDB 12.0 version (18)

Finally, we analyze the contribution of different sources (i.e., classic, HT, and/or computational methods) by type and by category, to the confidence level of the collection of RIs.

## 2 Results

A regulatory interaction is one of the major concepts, together with transcription units and operons, that describe the knowledge of regulation of transcription initiation. As with any piece of knowledge, it can be described at different levels of detail; at a low level we can say it is the triplet formed by the TF, the target gene, and the positive or negative effect. The basic requisites to annotate a new RI are one evidence of a TF binding near the gene start and evidence showing that the presence/absence of this TF has an effect positive or negative over the transcribed gene, that we call the evidence of function. The whole transcriptional regulatory network (TRN) at this high level is available as a downloadable file at https://regulondb.ccg.unam.mx/datasets, in the network interactions file “NetworkRegulatorGene.” Note that all other datasets within the group of “Regulatory Network Interactions” entail this low level of detail.

In contrast, at a high level of detail, knowledge of RIs involves the TF, the effector that affects its binding and unbinding conformation, the precise TF regulatory binding site, the regulated promoter, and the effect of the TF when bound, either activating or repressing transcription initiation. It may also be expanded and include knowledge of the TU that the promoter transcribes and therefore the set of regulated genes, and finally the growing conditions (experimental and control) where such regulation takes place. These fundamental concepts originated in the 1960s in the work of Jacob and Monod with the emergence of molecular biology in microbial organisms (19-21). Since then, many regulatory systems have been dissected and their molecular components identified. A group of experts, including one of us, has recently updated these concepts given the huge expansion of knowledge (22). We did some modifications, both in RegulonDB and EcoCyc, to conform to these new proposals. One that is relevant to this work is the distinction between TF binding sites (TFBSs) and the subset of TF regulatory sites (TFRSs), which are those TFBSs with functional evidence showing they have a regulatory role. This distinction is particularly relevant since genomic public data for *E. coli* is currently dominated by methods that identify TFBSs; only a few of them have evidence of their regulatory role on target genes or promoters, while most of them lack differential expression of the target genes. Fortunately, in RegulonDB we have incorporated the distinct notation of TFBSs and TFRSs, following (22). TFBSs *per se* do not support RIs.

Incidentally, the current confidence level for RIs is limited to their TF site binding evidence. This is no surprise; certainly, our curation and most knowledge provides evidence supporting genetic elements, with little or close to zero evidence codes and confidence levels for the interactions among these objects.

### 2.1 Updated and new evidence codes

As mentioned before, we wanted to distinguish classic vs HT methods and increase their precision to match with specific methods. We updated our table of evidence types, and we have modified their descriptions to explicitly include whether they are experimental methods, either HT or classic methods, or nonexperimental, such as computational predictions or author statements. We modified the names of evidence codes to make them more informative. Since some objects are rather complex, particularly the RIs, we have separated the evidence for binding within sites and the evidence for function within the RI itself. This also facilitates user searches for specific references whenever they come from different publications. Each evidence type is associated with a specific code, which we created intentionally keeping it as short as possible but informative and with prefixes indicating if it is an HT method.

We added a link in RegulonDB that offers the name, description, evidence code, and confidence level (see below) of all evidence types, as well as whether they correspond to *in vitro* or *in vivo* binding experiments. **See:** https://regulondb.ccg.unam.mx/manual/help/evidenceclassification. Although this table shows updated codes for RIs, promoters, and TUs, we focus in this paper only on the work around RIs. As can be seen, the new evidence types added are essentially those that support HT methods.

### 2.2 Confidence derived from multiple independent methods

Years ago, we classified in RegulonDB the different evidence types into either weak or strong, depending on the confidence that the methods provided to support the existence of a piece of knowledge. The general principle was that strong confidence comes from experiments that provide clear physical evidence of the existence of the object. For instance, binding of purified proteins in the case of a given TF binding to its binding site is considered strong evidence, whereas binding of cellular extracts is considered weak evidence. A limitation to this initial approach was that even if some objects are identified by different methods, either in the same paper or through the years in more than one publication, we did not have a process to add multiple weak evidence types and consider it a strongly supported object. This is contrary to a fundamental strategy in natural science, whereby further support to knowledge is gained by different, and ideally independent strategies or methods. We analyzed which of the different methods can be considered independent because they use different assumptions and/or different methodological strategies such that their potential sources of error are different (23). It is on this basis that we built our algebra to combine multiple weak independent sources of methods into a strong confidence level. We also proposed the combination of independent strong evidence types to create the new “confirmed” level of confidence.

In the current update, we have kept the same principles and criteria as defined in the 2013 paper and updated the three levels of confidence, given the increase in evidence types. In the case of classical evidence, data come from individual experiments focused on individual objects, so they were classified as strong, except the binding of cellular extracts, which can be considered less specific, because such experiments do not eliminate the possibility of indirect effects. HT binding evidence types were classified as weak, since they involve several processing steps, including different bioinformatics options of methods and thresholds, making the final results more variable and dependent on the specific set of programs and variables used in their final identification. Thus, processing the same raw data could potentially result in different final collections of objects; in addition, there is no consensus yet on a uniform processing pipeline used by the community. Nonexperimental evidence types were also classified as weak; however, among them only computational analysis can be used in combination with other evidence types to upgrade the confidence level of the RIs associated, while author statements or inferences by curators do not allow such an upgrade. As a result, our current types of evidence for RIs, their classification in groups, and their levels of confidence are summarized in Figure 1.

**Figure 1.**
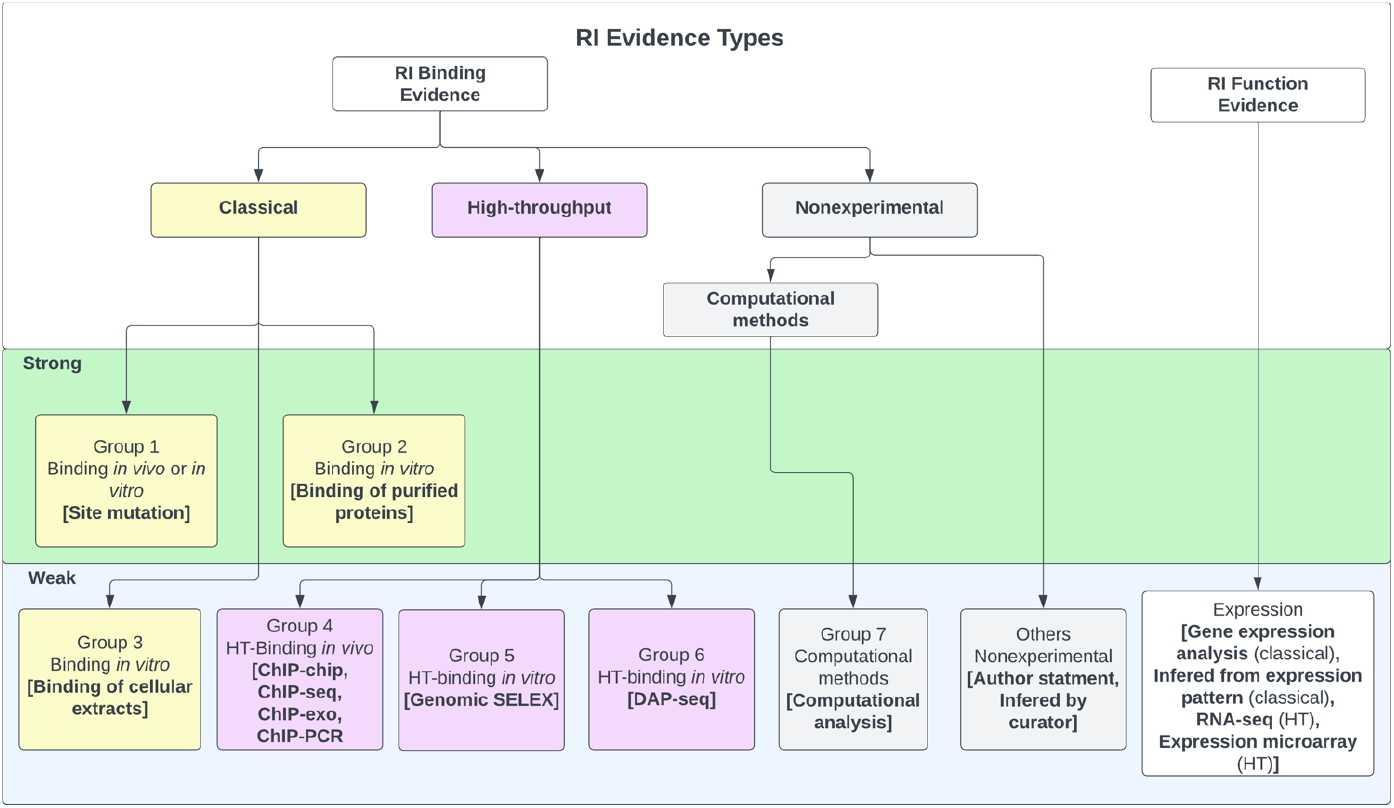
Evidence types for RIs. Any RI requires evidence for the binding of the TF, together with functional evidence showing its regulatory effect in transcriptional activity. Evidence types are grouped in three major categories (classical, HT and nonexperimental), each specific group contains methods that are not considered independent, and methods of different groups are considered independent evidence. The algebra of their independent groupings is limited to binding evidence types, which define the level of confidence as discussed in the main text.

We assigned each evidence type to one of seven possible groups (Figure 1) and defined the combinations that upgraded the object confidence level using those group numbers. Evidence types in the same group are considered to share methodological bias and cannot be combined to upgrade the confidence level of the associated object, while evidence types between different groups are considered independent and their combination can upgrade the object confidence level. Currently, only the evidence types of groups 1 and 2 are classified as strong, groups 4 to 8 are considered weak evidence types, and the group of “others” do not contribute to confidence (Figure 1). The evidence types 4, 5, and 8 belong to the category HT; the evidence types from group 4 are considered independent from the 5 and 8 types because the methods are considered different enough, with the first group assayed *in vivo* while methods of groups 5 and 8 are identified binding *in vitro*; the experimental and computational processing of raw data are also different. gSELEX and DAP-seq are also considered independent of each other. Based on these groupings, the different evidence for an individual RI can be combined to increase its confidence level as follows, remembering that any RI must have functional evidence:

1. Two independent binding evidence types with confidence level “strong” (groups 1 or 2) upgrade the object confidence level to “confirmed”.
2. Two independent binding evidence types with confidence level “weak” (groups 4 to 7) upgrade the object confidence level to “strong”.
3. Two independent binding evidence types with a confidence level of “weak” (groups 4 to 7) in addition to a strong evidence type (group 1 or 2) upgrades the object confidence level to “confirmed”.
4. Four independent binding evidence types with a weak confidence level (groups 4 to 7) upgrades the object confidence level to be confirmed.

It is worth mentioning that two independent weak evidence types can upgrade the object evidence to strong only when the evidence for the effect in regulation is not missing in RegulonDB. Note that binding of purified protein and site mutation are currently the only evidence types with confidence level strong. There is no single evidence that supports the level of “confirmed”. Site mutation is classified as a strong evidence type because it involves the precise identification of the regulatory site and TF binding since, if modified, even in the presence of the TF, there is no effect on transcription, either *in vivo* (24,25), through a reporter gene or by *in vitro* transcription in the presence or absence of a determined TF (26). Binding of purified protein includes two similar methodologies: electrophoretic mobility shift analysis (EMSA) and footprinting, in which the TF binding to a specific sequence target is probed *in vitro*. Note that currently only HT methods are sufficient to provide a confirmed confidence level, as there are three independent HT groups of methods, and in addition some HT methods (i.e. ChIP-seq) frequently add a computational identification of the binding site enhancing its confidence level from weak to strong.

The complete set of evidence type combinations that upgrade an RI confidence level can be found under the “regulatory interactions” of the “Stage II. Assignment of confidence level based on additive evidence types” section of the webpage https://regulondb.ccg.unam.mx/manual/help/evidenceclassification. For instance, ChIP-chip, ChIP-seq, and ChIP-exo belong to group 4, whereas gSELEX belongs to group 5. The rule (4/5/3)-S means that if an RI has evidence from any method in group 4 plus any evidence from group 5, together they upgrade two weak binding evidence types into a strong confidence level. Remember that the evidence of the regulatory effect is always required for an RI.

Once all these updates were in place, we recalculated the confidence levels for the two versions of the complete set of RIs present in RegulonDB, i.e., the version before and the one after adding the binding evidence of all binding HT collections. This is presented in the section on the “Anatomy of Knowledge.” Before that discussion, we explain another implementation that enhances the quality of knowledge representation of RIs in RegulonDB.

### 2.3 Three representations of regulatory interactions: TF-promoter, TF-TU, and TF-gene

A different challenge we have addressed when searching for the best possible way to reflect knowledge is the need for intelligent ways to deal with partial knowledge. It is not uncommon for a curator to have to choose the least costly assumption when knowledge is lacking. For instance, years ago, since by definition a TU has a promoter, we added the so-called “phantom promoters” to those TUs that had no characterized promoter. This was eventually eliminated as suggested by Rick Gourse in an EcoCyc meeting, to avoid confusion by users. Another example illustrating the same problem was how to deal with the curation of RIs. Historically, we curated RIs affecting a given promoter, even when there was no such specific evidence. The curator uploaded the RI when the target gene had only one promoter, and if the target gene had two or more promoters, the new RI was mentioned in notes of the TU. It is important to be aware that our curation work has been evolving for more than 30 years now. In this long period, we have added new objects, new features, improved our evidence codes, in addition to many more changes, essentially improving the quality of knowledge representation.

In order to minimize assumptions in our curation process, we defined three levels of description of RIs, which we use depending on the level of detail of knowledge available. We call these “RI types”:

1. The most precise knowledge is when there is evidence that identifies the regulated promoter affected by an RI. Most of these come from classical experiments. In these cases, it is reasonable to deduce that TUs associated with the regulated promoter are regulated by the new RI. These are RIs described at the level of “TF-promoter.”
2. A less detailed description is when the regulated promoter is not known and there is evidence of a change in expression of a group of adjacent genes on the same strand of the promoter that matches with an existing TU with or without promoter. In such cases, we associate the new RI to the existing TU. We call these TF-TU RIs. If there is no previous TU, we create a new TU without a promoter and with evidence of coexpression and link to it the new RI.
3. Finally, when the regulated promoter has not been identified and there is evidence of differentially regulated transcription of the downstream gene(s) from a TF binding site, we create a new RI for which the target is the gene. We call these TF-gene RIs.

As a result, we currently have three means of adding RIs, depending on available knowledge in RegulonDB and EcoCyc: TF-promoter, TF-TU, and TF-gene.

The curation of knowledge related to RIs exerted by a TF depends on several rules. The easy case is when there is not a previously annotated RI with the same TF and gene; in this case, a new RI at the adequate level is annotated, according to the available knowledge. However, if there is a previous RI, of any of the three types, and the new and previous knowledge match, the new evidence is added to the existing RI. This involves an “RI mapping” process (See methods).

As a result of all the modifications described in section II, we have made public RegulonDB version 12.0, which includes the updated collections of RIs, either separated as TF-promoter, TF-TU, or TF-gene. The RI set has each RI with its complete list of evidence types, enabling users to exclude for instance, ChIP-seq evidence and recalculate an improved gold standard for new ChIP-seq experiments that prevents evaluating ChIP-seq data with previously performed ChIP-seq data. Users can access in the downloadable files each type of RI, or the union of all of them.

### 2.4 Incorporation of HT-binding evidence to the existing RIs

We have been systematically curating RIs for HT published data. It is important to note that until now, HT-supported RIs were identified only when evidence of binding and function were reported in the same publication. Certainly, most studies reporting genome-wide TF binding do not report TF-dependent differential gene expression; in some cases, it is assayed for a small set of TF-binding target genes. An interesting alternative to maximize the use of this data is to map the peaks from the HT-binding datasets to existing RIs in RegulonDB and add such HT-binding evidence to known RIs. We performed this process, enriching the evidence and increasing the confidence levels for existing RIs, although many RIs from the HT datasets await their functional evidence.

The current total number of RIs in RegulonDB is 5,466 from 237 different TFs of which 148 have at least one HT dataset, 27 putative TFs have an HT-TFBSs dataset but have no RIs in RegulonDB. The enrichment of binding evidence for 1329 RIs resulted in changes in the RI confidence levels as well as in the evidence categories from “nonexperimental” to “HT” and from “classical” to “classical & HT” as discussed below (see Tables 1 and S1).

**Table 1.**
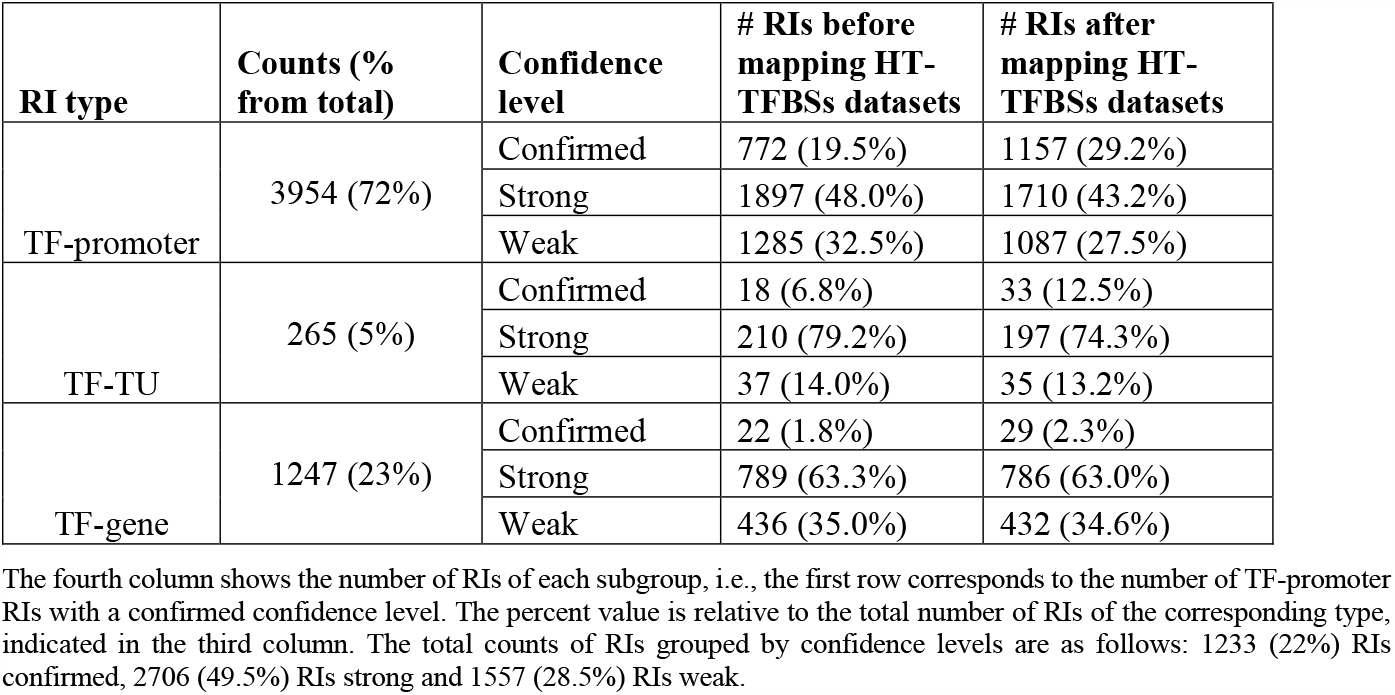
Number of RIs grouped by type and by confidence level.

### 2.5 Anatomy of knowledge supporting the RIs

Given the improved level of detailed updated annotations for all RIs currently available in RegulonDB, we used them to analyze the internal anatomy of this corpus of knowledge (Tables 1 and S1). We could expect, for instance, that RIs of the TF-promoter type come mostly from classical methods, and these probably include the most confirmed interactions. In this section, we show how the data helped us to answer these kinds of questions.

#### 2.5.1 Classical evidence dominates TF-promoter interactions with confirmed and strong confidence levels

First, we wanted to see if RIs of the three different types contribute differently to the confidence level. Our analysis showed that 72% of RIs belong to the type TF-promoter, 23% are TF-gene, and only 5% are TF-TU (Table 1, Figure 2A). In terms of confidence levels, currently, RegulonDB (version 12.1) contains a total of 5,466 RIs, of which 1,219 (22.3%) have confirmed evidence, 2,693 (49.3%) have strong evidence, and 1,554 (28.4%) have weak support (Table 1). Combining the confirmed and strong levels includes almost 70% of all current RIs.

**Figure 2.**
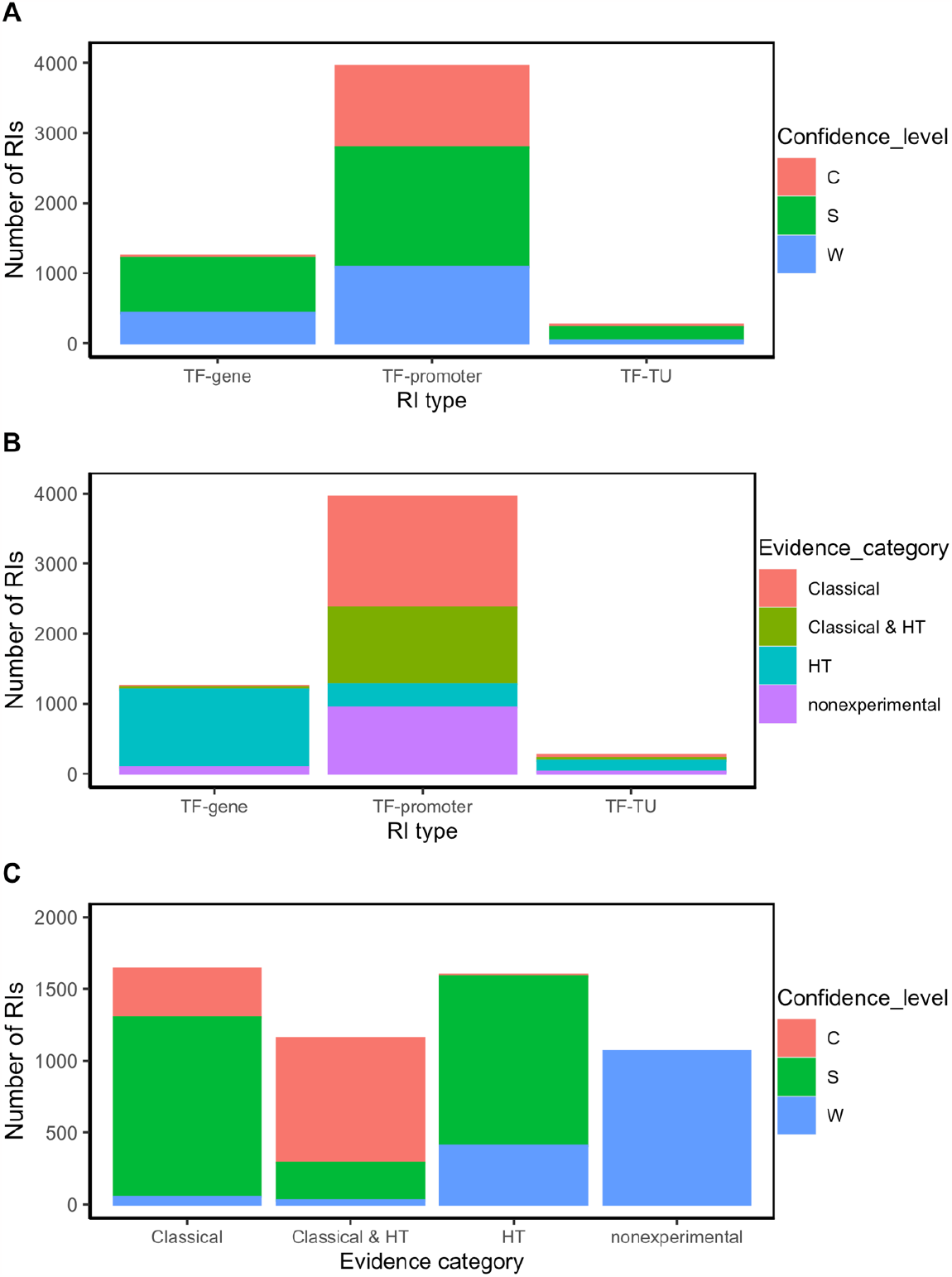
RI distribution analysis by type of RI (TF-promoter, TF-TU or TF-gene), confidence level (C: confirmed, S: strong, and W: weak), and evidence category (classical, HT and nonexperimental) A) Number of RIs by confidence level for each type of RI; B) Number of RIs by evidence category for each type of RI. C) Number of RIs by confidence level for each evidence category.

We have classified the binding evidence types supporting RIs (and other objects in RegulonDB) in three general categories, classical, HT and nonexperimental as shown in Figure 1. RIs can be supported by combinations of evidence types belonging to different categories, so for our analysis we assigned the global categories: “classical”, “HT”, “classical & HT” and “nonexperimental”, the first three can include or not nonexperimental evidence. For these global categories there are 1640, 1601, 1157 and 1068 RIs, respectively. RIs with classical evidence (classical + classical & HT) represent a 51.2% (2797) and most likely this fraction will diminish with time.

As expected, the TF-promoter type is the one with the highest number of strong and confirmed levels of confidence (Figure 2A); this is no surprise since, as mentioned before, the TF-promoter level is the one where more mechanistic knowledge of the RI is known. To date, 67.6% (2674) of TF-promoter RIs have been characterized by classical methods (Figure 2B, Table S1).

TF-gene and TF-TU RIs are mainly supported by HT evidence (Figure 2B), and they have mostly strong or weak confidence levels (Figure 2A).

The most frequent combination supporting confirmed RIs is “site mutation” (classical evidence) with “binding of purified protein” (classical evidence) and “computational analysis” (nonexperimental evidence) (Figure 3A), followed by “binding of purified protein” (classical evidence) combined with “genomic SELEX” (HT evidence) and “computational analysis” (nonexperimental evidence) (Figure 3A)

**Figure 3.**
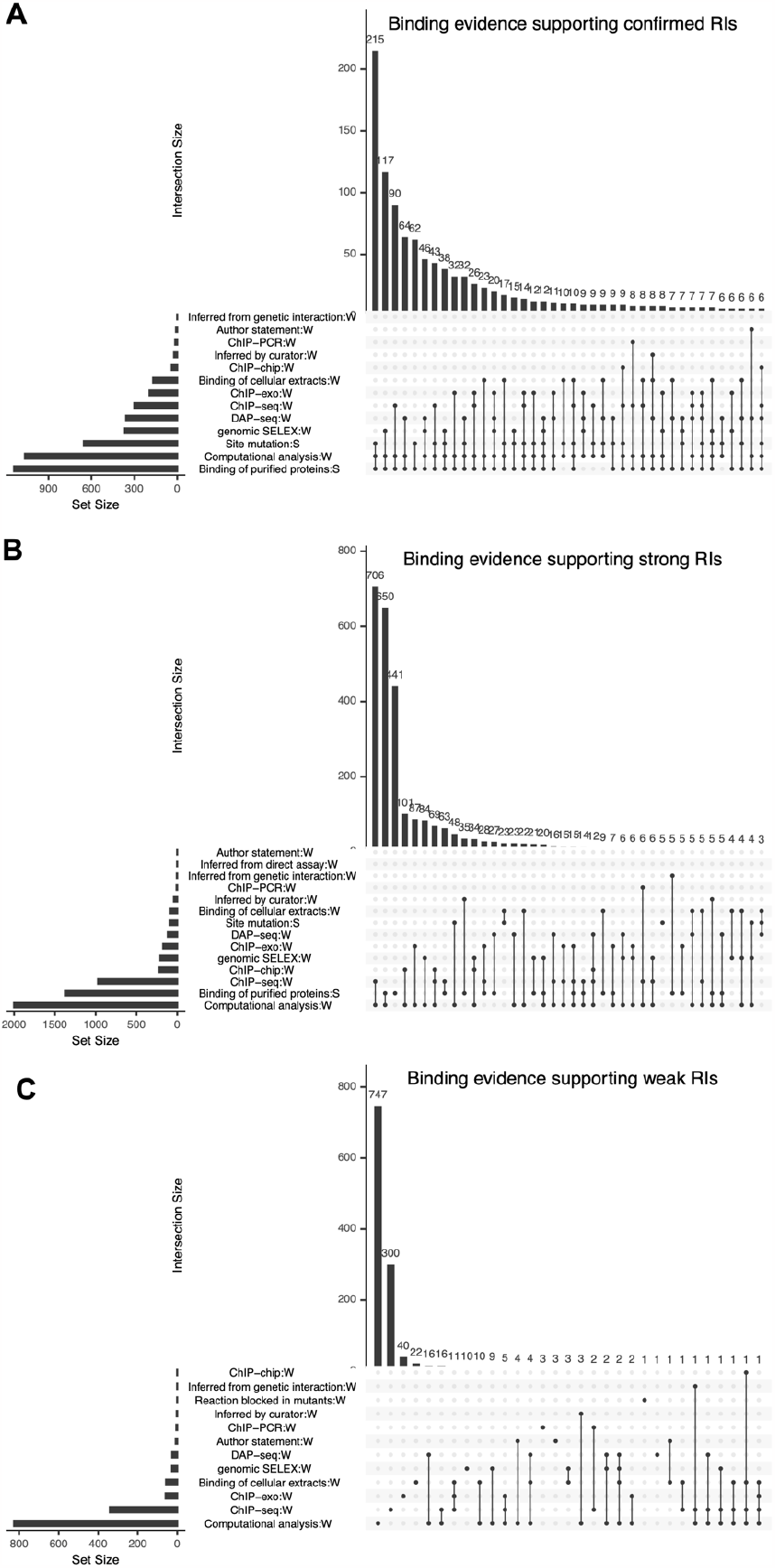
High detail combinations of binding evidence supporting RIs for each confidence level confirmed (A), strong (B), or weak (C). The number of RIs with each combination is shown on each bar. The Y-axis gives the number of RIs of intersections.

Several RIs supported by HT evidence, and without classical evidence, have strong confidence level. An interesting consequence of the integration of evidence from HT datasets is that now there are RIs at the confirmed level supported exclusively by HT evidence. In fact, the two most frequent combinations supporting strong RIs are: 1) “ChIP-seq” (HT evidence) combined with “computational analysis” (nonexperimental evidence) (Figure 3B), and 2) “binding of purified proteins” (classical evidence) combined with “computational analysis” (nonexperimental evidence) (Figure 3B).

#### 2.5.2 Most of the weak RIs are supported by nonexperimental or HT evidence

Most of the weak RIs are supported only by “computational analysis” or by “ChIP-seq” evidence types (Figure 3C). As mentioned before, HT evidence is considered weak, so RIs supported by only ChIP-seq have a weak confidence level (Figure 3C). There are also some weak RIs supported by classical evidence types, for example, RIs supported only by the evidence of “binding of cellular extracts”. In Figure 3C, we can observe that some RIs are supported by different combinations of independent evidence types; however, they do not become strong, because for these RIs the evidence of function (effect over expression) is missing, probably due to the historic process of curation. Future curation will enable us to recover their functional evidence. Note that 100% of nonexperimental RIs are classified as weak (Figure 2C).

#### 2.5.3 How HT evidence is changing the landscape of knowledge

Taking as reference the current set of RIs with all evidence types associated, if all HT-binding evidence were deleted, the confidence level would be considerably affected, with decreases of 44.5% for confirmed RIs and 24.6 % for the strong RIs (Figure 4).

**Figure 4.**
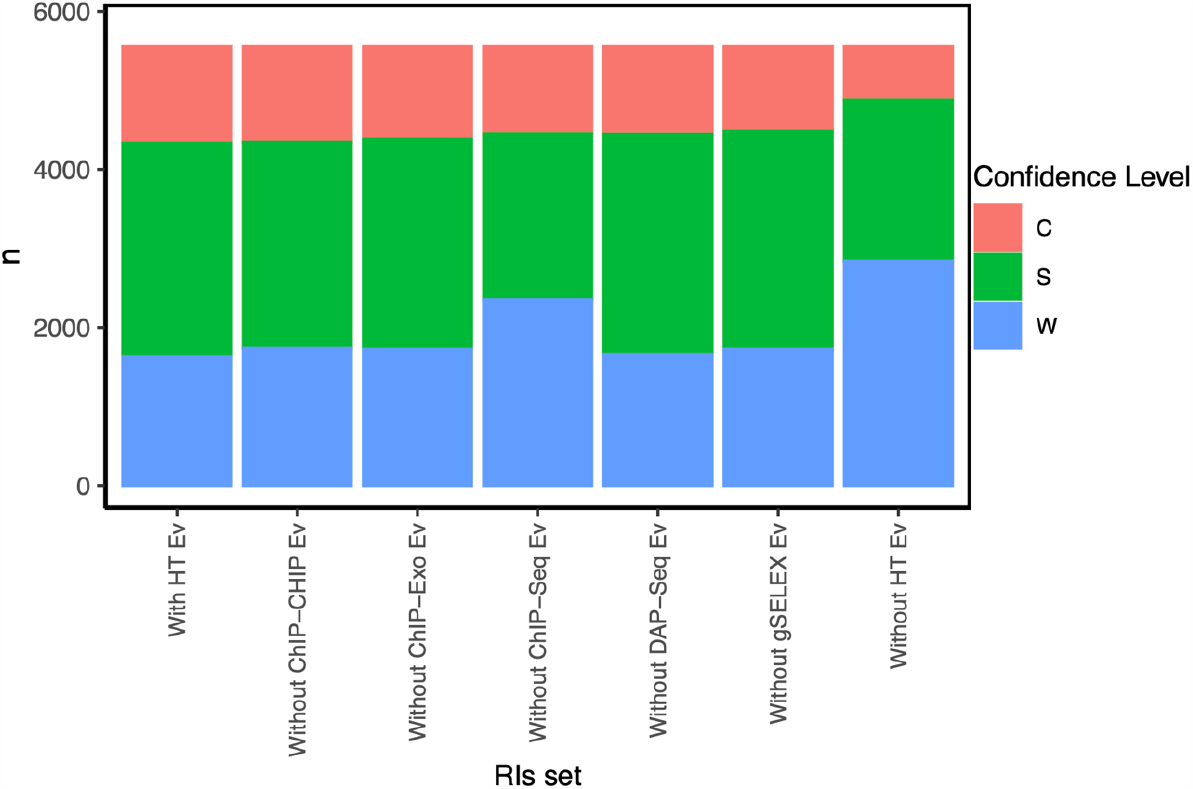
Confidence level from the RI set distribution, with and without HT-binding evidence.

We know that this contribution involves strengthening the evidence of RIs identified by classical methods and by the identification of new RIs through multiple HT methods, i.e., confirmed and strong RIs with classical and HT evidence and strong RIs with only HT evidence (Figure 2C). We found that currently, ChIP-seq is the methodology that contributes the most in increasing the number of strong RIs (Figure 4). This is no surprise given that this method currently contributes to the largest number by far of RIs.

#### 2.5.4 Use of gold standard datasets to benchmark HT-binding methodologies

The efforts that have been put into evidence curation for RIs using specific codes for different methodologies, along with their classifications into independent groups, confidence levels, and categories, now allow us to filter and create subsets of RIs that can be used as a gold standard for benchmarking HT-binding methodologies. The complete set of RIs is available on the RegulonDB website under “Releases & Downloads/Downloads/Experimental Datasets/TF-RISet”. On this page, users can download the entire set, and there are also two tools available:

1. Browse and Filter: In this tool, filters can be applied to each column to obtain a subset of RIs, and users can download them accordingly. For example, RIs with a confirmed confidence level could be filtered.
2. Confidence Level Calculator Tool: In this tool, one or multiple evidence codes can be selected to be ignored, and the confidence level can be recalculated.

As mentioned, the results of HT methodologies are frequently compared with the RegulonDB data as a way to validate them. However, these analyses had been performed with the complete set of RIs with all sorts of evidence supporting them. Now, specific gold standard datasets that exclude specific sources, to avoid circularity, can be used.

To assess the performance of HT-binding methodologies in recovering sites from classical RIs in RegulonDB, we considered only the subset of TFs that have at least one classical RI. For each TF we calculated the percentage of classical RIs that map with the peaks in the corresponding dataset, the average percentage was calculated for the subset of TFs of each HT methodology. These analyses include RIs belonging to all three confidence levels: weak, strong, and confirmed. The results are depicted in Figure 5. ChIP-seq was the methodology that recovered the highest percentage (76.8 +/-20.4%) of classical RIs at site level, followed by gSELEX (65.9 +/-34.5%), DAP-seq (51.0 +/-35.6%) and ChIP-exo (33.1 +/-38.5%). A different form of compare is to calculate percentage of the total number of classical RIs mapped for the subset of corresponding TFs. Using this approximation, similar results are obtained with 70.1% of total RIs recovered by ChIP-seq, 51.3% recovered by gSELEX, 28.0% recovered by DAP-seq and 22.6% recovered by ChIP-exo (Supplementary_material_2).

**Figure 5.**
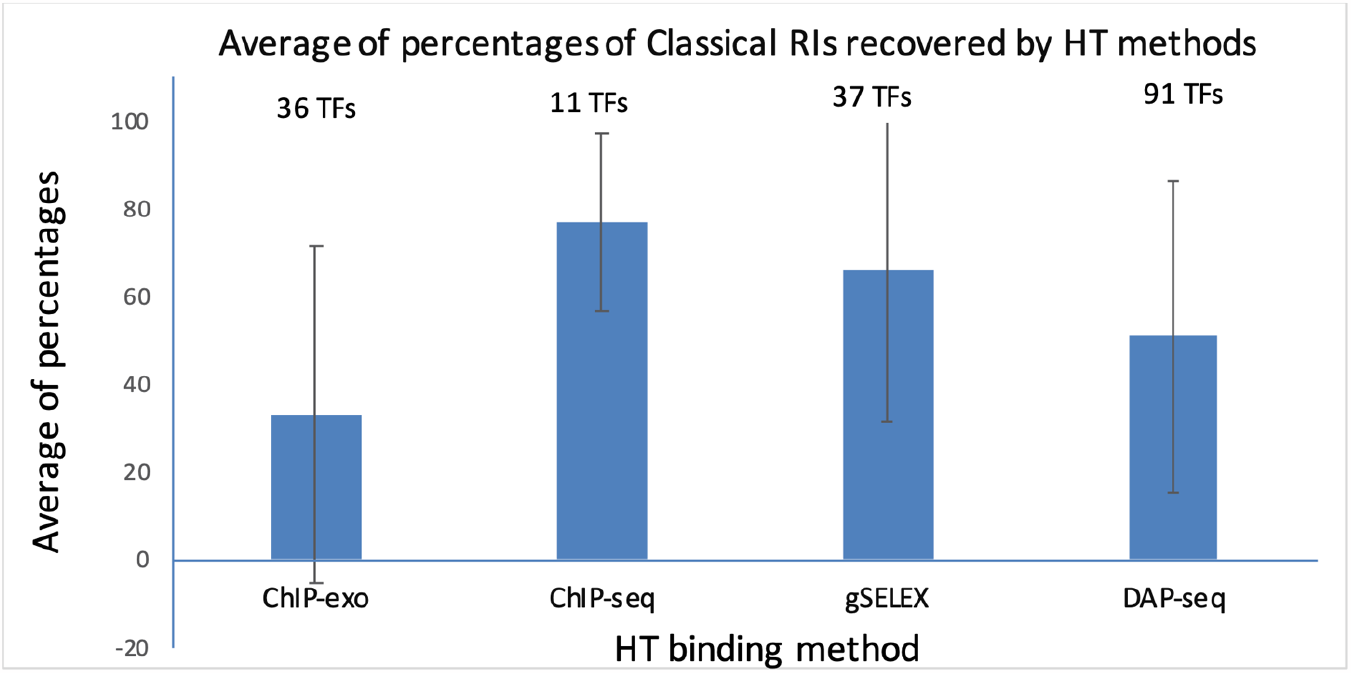
Comparison of recovered classical RIs by different HT-binding methodologies. For each methodology, the fraction of recovered RI sites in RegulonDB was estimated and the average for all TFs and std deviation is shown. The set of TFs is specific to each method given the currently available datasets gathered in RegulonDB version 12.1 and also limited to those TFs for which there is at least one classical RI in RegulonDB (For data details see Suppementary_material_2).

Subsequently, to be stricter, the same analysis was performed using the dataset of RIs with a confirmed confidence level without considering HT evidence. As expected, this subset includes only RIs with at least one classical type of evidence for binding, the results are depicted in Figure S2. Once again, ChIP-seq was the methodology that recovered the highest average percentage (95.5% +/-7%) of classical confirmed RIs, followed by gSELEX (77.6% +/-35.0%), DAP-seq (57.5% +/-39.7%) and ChIP-exo (35.4% +/-39.1%) (Supplementary_material_3).

## 3 Discussion

One relevant outcome of this work is the availability of gold standard datasets useful for benchmarking new methodologies. From the master RI complete table (“Releases & Downloads/Downloads/Experimental Datasets/TF-RISet”) containing all the evidence types for each RI, users can make their own combinations. Users can include or exclude specific subcollections based on the method and/or evidence types and can also select subsets of RIs by filtering by confidence levels with the tools described before.

As shown, we performed a series of improvements, including more precise evidence codes, the three types of RIs adequate to capture the diverse cases of partial knowledge, and the updated calculation of confidence levels of RIs. These advances enable the analyses performed of the different sources of knowledge and their contribution to the currently known *E. coli* TRN. Figure 4 shows how the set of RIs increases in confidence levels when the HT binding evidence is considered. It is clearly an illustration of how in the near future HT methods will most likely dominate and expand the knowledge supporting the *E. coli* TRN. Although the comparison is limited to RIs, the same change in the anatomy of knowledge sources will also include other elements such as TSSs and TUs.

We used as the gold standard the set of RIs with at least one classical evidence and with a confidence level confirmed to compare the datasets from the different HT methodologies in our current collection. It is true that this is a preliminary and incomplete comparison given the limited data. Certainly, the number of datasets and of TFs tested is quite different for each method, the laboratories are also different, briefly, these datasets were not generated within a pre-planned strategy for a well-defined comparison. Within these limitations, we can see that the ChIP-seq collection, coming from different laboratories, shows a significantly higher fraction of recovered sites from RegulonDB compared to the other three methods. The comparisons remain consistent in the three different ways of averaging the results as shown in Figures 5 and S2. This is surprising since as shown before, the largest fraction of RIs in RegulonDB is still coming from *in vitro* classical methods, whereas ChIP-seq is an *in vivo* methodology.

A set of gold standard data is useful for different fields of biomedical research, but it must be not only a reference collection but also one that represents data with the highest level of confidence. The evaluation of data confidence based on independent evidence is commonly used in specific investigations; for example, quantitative reverse transcription PCR (RT-qPCR) is used to validate RNA-seq experiments. However, only a few studies have used this approximation to evaluate data on a large scale (27). In medicine, levels of evidence are assigned to studies based on diverse criteria, such as quality, with higher levels of quality of evidence entailing less risk of bias (28). Our approach can be applied to analyze data from curated databases which have structured evidence codes associated with objects, such as BioGRID database (29), or that can be applied to other cellular processes from *E. coli* to determine, for example, which are the best-characterized metabolic pathways, based on data from EcoCyc.

The different improvements discussed in this paper enable us to incorporate HT-generated knowledge without “diluting” the valuable fraction of knowledge supported by classical molecular biology methods, since it is easy to dissect subsets based on their supporting methods. As shown with RIs in this paper, we will move to adequately combine classical and HT methods for TSSs and TUs and update what constitutes one of the best-characterized TRNs of any microbial organisms, which is also likely the best computationally represented corpus of knowledge of gene regulation.

## Methods

### 3.1 Updates in RegulonDB

#### Evidence update

In RegulonDB version 12.1, we made important evidence-related changes, including: 1) Evidence code. The evidence codes were made more informative, i.e., BPP was changed to EXP-IDA-BINDING-OF-PURIFIED-PROTEINS. 2) Evidence confidence levels. Evidence types were classified as “weak” or “strong” depending on whether they provided physical and direct proof of the existence of the object or interaction

#### Object confidence level update

The confidence level for each RI, promoter, and transcription units was calculated and updated using the linked evidence and the additive evidence. The confidence level assignment to RIs is described below; we followed the same principles for the other objects. These changes are reflected in the RegulonDB interface as well as in the downloadable datasets.

### 3.2 Input for RIs

The analysis was done using RegulonDB version 12.1 synchronized with Ecocyc version 27.0. In this version, the downloadable text file for Regulatory Interactions was made available and also the evidence catalog file. The formats and descriptions of these files are available at https://github.com/regulondbunam/download-data-files.

### 3.3 Confidence level assignments to evidence types and to RIs

The confidence levels were assigned to RIs by a process involving two stages:

**Stage I**. Each single evidence type was classified into weak or strong, as described in section 2.2.

**Stage II**. Assignments of confidence levels to RIs were based on the set of their evidence types.

In this stage, we use the concept “additive evidence,” which in previous versions was called “cross-validation.” As we proposed a while ago, Weiss et al. (2013) (23), the confidence level of a biological entity depends on the combined evidence derived from mutually independent methods.

We grouped methods that could have similar sources of false positives. This resulted in seven independent evidence groups (Figure 1). The combinations of evidence groups that upgraded the RI confidence levels were defined based on the four rules mentioned in section II.2. We call these combinations additive evidence, which define the final level of confidence assigned to each RI. The complete set of group combinations that upgraded RI confidence levels can be found under the “regulatory interactions” of the “Stage II” section of the webpage: https://regulondb.ccg.unam.mx/manual/help/evidenceclassification.

### 3.4 Access options for users

Although it is well known that RegulonDB contains the comprehensive collection of experiments performed through decades of classic methodologies, users must be aware that we already have incorporated evidence from HT methods.

The current publicly available RegulonDB offers downloadable datasets grouping collections of objects in https://regulondb.ccg.unam.mx/datasets. The first option offers the “RIset” which contains all the evidence types for binding and function of RIs, in columns 21 and 22 respectively. These can be used to filter and extract, for instance, the subcollection supported only by classic methods. The same strategy could be used to select RIs supported by a specific evidence type. Users can also subselect RIs based on the confidence level, or on the different groups of methods, as described in Figure 1. Furthermore, users may define their own rules and categories of different levels of confidence and use the whole collection of evidence types to classify each individual RI in a new classification of confidence levels.

All scripts and computational processes built to generate the data and analyses presented in this paper are publicly available and can be found at https://github.com/PGC-CCG/supplementary-material/tree/master/gold-standard

### 3.5 Analysis of the current set of RIs

The analyses of the anatomy of RI knowledge presented here were performed using R (2022.06.23, version 4.2.1), Rstudio (2022.07.1, Build 554), and the ggplot2 (version 3.4.0) library.

### 3.6 Mapping collection of TFBSs-HT to TFRSs from RegulonDB

The collection of HT TFBSs contains four subcollections: DAP-seq, ChIP-seq, ChIP-exo, and gSELEX (1). In order to make these collections comparable among them and with RegulonDB TFRSs, multiple steps were implemented that together constituted what we call “mapping,” in this case mapping of HT-binding data with known sites. This mapping involves:

#### 3.6.1 Uniformization of the genome coordinates for all datasets

The coordinates of the DAP-seq datasets were published using the last genome version of the *E. coli* str. K-12 substr. MG1655 (U00096.3), so they were not modified. The ChIP-seq, gSELEX, and ChIP-exo datasets with coordinates in the past genome version (U00096.2) were updated to version U00096.3. The corresponding Scripts are found in the github indicated

#### 3.6.2 To map the RegulonDB RI set with peaks from the HT-TFBSs subcollections

A program in Python was implemented that compared each RI binding site with each peak corresponding to the same TF. A match is assumed when the RI site coordinate is within the region covered by the HT peak.

When a match between an RI and the HT data is found, the evidence of the corresponding HT-methods is added to the corresponding RI. This process is executed in each RegulonDB release. Scripts are found in the github as mentioned.

### 3.7 HT-binding methodology efficacies in recovering sites from classical RIs in RegulonDB

For the comparison of ChIP-seq, ChIP-exo, gSELEX, and DAP-seq, the RIs set was mapped to the complete collection as described before. The fraction of RIs with at least one piece of classical evidence that were recovered by each method for each TF was then calculated. TFs in each collection with zero RIs featuring at least one classical evidence were excluded from this analysis. The same algorithm was applied to determine the proportion of RIs with confirmed confidence levels without considering HT evidence in the calculation.

### 3.8 Resource Identification Initiative

To take part in the Resource Identification Initiative, please use the corresponding catalog number and RRID in your current manuscript. For more information about the project and for steps on how to search for an RRID, please click <u>here</u>.

## 5 Author Contributions

P.L. Curation, redefinition of evidence codes and their additive groups; coordination with the computational team and their developers; software for mapping and data analysis, discussion, and writing. S.G.C.: Curation, redefinition of evidence codes and their additive groups, analysis and discussion. H.S.: Software, validation, visualization, review and analysis. C.R. Validation, analysis, discussion. V.H.T. redefinition of evidence codes, and their additive groups; curation of DAP-seq datasets. L.M.R.: Software for additive evidence, release process and downloading datasets, C.B.M: Software for normalization, and genome coordinates updates; J.C.V.: Conceptualization, supervision, leading the research process, funding acquisition, review and writing.

## 7 Acknowledgments

We acknowledge Gabriel Alarcón-Carranza, Felipe Betancourt-Figueroa and Andrés G. López-Almanzo for the design and implementation of the Confidence Level Calculator Tool and the updates of evidence and related data in RegulonDB. We also acknowledge IT support by Víctor Del Moral.

## References

1. Tierrafría, V.H., Rioualen, C., Salgado, H., Lara, P., Gama-Castro, S., Lally, P., Gómez-Romero, L., Peña-Loredo, P., López-Almazo, A.G., Alarcón-Carranza, G. et al. (2022) RegulonDB 11.0: Comprehensive high-throughput datasets on transcriptional regulation in Escherichia coli K-12. Microb Genom, 8, PMID: 35584008

2. Kahramanoglou, C., Seshasayee, A.S., Prieto, A.I., Ibberson, D., Schmidt, S., Zimmermann, J., Benes, V., Fraser, G.M. and Luscombe, N.M. (2011) Direct and indirect effects of H-NS and Fis on global gene expression control in Escherichia coli. Nucleic Acids Res, 39, 2073–2091, PMID: 21097887

3. Shimada, T., Yamamoto, K. and Ishihama, A. (2011) Novel members of the Cra regulon involved in carbon metabolism in Escherichia coli. J Bacteriol, 193, 649–659, PMID: 21115656

4. Shimada, T., Fujita, N., Yamamoto, K. and Ishihama, A. (2011) Novel roles of cAMP receptor protein (CRP) in regulation of transport and metabolism of carbon sources. PLoS One, 6, e20081, PMID: 21673794

5. Shimada, T., Bridier, A., Briandet, R. and Ishihama, A. (2011) Novel roles of LeuO in transcription regulation of E. coli genome: antagonistic interplay with the universal silencer H-NS. Mol Microbiol, 82, 378–397, PMID: 21883529

6. Seo, S.W., Kim, D., Latif, H., O’Brien, E.J., Szubin, R. and Palsson, B.O. (2014) Deciphering Fur transcriptional regulatory network highlights its complex role beyond iron metabolism in Escherichia coli. Nat Commun, 5, 4910, PMID: 25222563

7. Shimada, T., Takada, H., Yamamoto, K. and Ishihama, A. (2015) Expanded roles of two-component response regulator OmpR in Escherichia coli: genomic SELEX search for novel regulation targets. Genes Cells, 20, 915–931, PMID: 26332955

8. Ishihama, A., Shimada, T. and Yamazaki, Y. (2016) Transcription profile of Escherichia coli: genomic SELEX search for regulatory targets of transcription factors. Nucleic Acids Res, 44, 2058–2074, PMID: 26843427

9. Shimada, T., Saito, N., Maeda, M., Tanaka, K. and Ishihama, A. (2015) Expanded roles of leucine-responsive regulatory protein in transcription regulation of the Escherichia coli genome: Genomic SELEX screening of the regulation targets. Microb Genom, 1, e000001, PMID: 28348809

10. Kim, D., Seo, S.W., Gao, Y., Nam, H., Guzman, G.I., Cho, B.K. and Palsson, B.O. (2018) Systems assessment of transcriptional regulation on central carbon metabolism by Cra and CRP. Nucleic Acids Res, 46, 2901–2917, PMID: 29394395

11. Shimada, T., Ogasawara, H. and Ishihama, A. (2018) Single-target regulators form a minor group of transcription factors in Escherichia coli K-12. Nucleic Acids Res, 46, 3921–3936, PMID: 29529243

12. Kroner, G.M., Wolfe, M.B. and Freddolino, P.L. (2019) Escherichia coli Lrp Regulates One-Third of the Genome via Direct, Cooperative, and Indirect Routes. J Bacteriol, 201, PMID: 30420454

13. Anzai, T., Imamura, S., Ishihama, A. and Shimada, T. (2020) Expanded roles of pyruvate-sensing PdhR in transcription regulation of the Escherichia coli K-12 genome: fatty acid catabolism and cell motility. Microb Genom, 6, PMID: 32975502

14. Choudhary, K.S., Kleinmanns, J.A., Decker, K., Sastry, A.V., Gao, Y., Szubin, R., Seif, Y. and Palsson, B.O. (2020) Elucidation of Regulatory Modes for Five Two-Component Systems in Escherichia coli Reveals Novel Relationships. mSystems, 5, PMID: 33172971

15. Ishihama, A. and Shimada, T. (2021) Hierarchy of transcription factor network in Escherichia coli K-12: H-NS-mediated silencing and Anti-silencing by global regulators. FEMS Microbiol Rev, 45, PMID: 34196371

16. Shimada, T., Furuhata, S. and Ishihama, A. (2021) Whole set of constitutive promoters for RpoN sigma factor and the regulatory role of its enhancer protein NtrC in Escherichia coli K-12. Microb Genom, 7, PMID: 34787538

17. Baumgart, L.A., Lee, J.E., Salamov, A., Dilworth, D.J., Na, H., Mingay, M., Blow, M.J., Zhang, Y., Yoshinaga, Y., Daum, C.G. et al. (2021) Persistence and plasticity in bacterial gene regulation. Nat Methods, 18, 1499–1505, PMID: 34824476

18. Salgado, H., Gama-Castro, S., Lara, P., Mejia-Almonte, C., Alarcón-Carranza, G., López-Almazo, A.G., Betancourt-Figueroa, F., Peña-Loredo, P., Alquicira-Hernández, S., Ledezma-Tejeida, D. et al. (2023) RegulonDB v12.0: a comprehensive resource of transcriptional regulation in E. coli K-12. Nucleic Acids Res, PMID: 37971353;

19. Colvill, A.J., Kanner, L.C., Tocchini-Valentini, G.P., Sarnat, M.T. and Geiduschek, E.P. (1965) Asymmetric RNA synthesis in vitro: heterologous DNA-enzyme systems; E. coli RNA polymerase. Proc Natl Acad Sci U S A, 53, 1140–1147, PMID: 4958034

20. Jacob, F. and Monod, J. (1961) Genetic regulatory mechanisms in the synthesis of proteins. J Mol Biol, 3, 318–356, PMID: 13718526

21. Chamberlin, M. and Berg, P. (1962) Deoxyribo ucleic acid-directed synthesis of ribonucleic acid by an enzyme from Escherichia coli. Proc Natl Acad Sci U S A, 48, 81–94, PMID: 13877961

22. Mejía-Almonte, C., Busby, S.J.W., Wade, J.T., van Helden, J., Arkin, A.P., Stormo, G.D., Eilbeck, K., Palsson, B.O., Galagan, J.E. and Collado-Vides, J. (2020) Redefining fundamental concepts of transcription initiation in bacteria. Nat Rev Genet, 21, 699–714, PMID: 32665585

23. Weiss, V., Medina-Rivera, A., Huerta, A.M., Santos-Zavaleta, A., Salgado, H., Morett, E. and Collado-Vides, J. (2013) Evidence classification of high-throughput protocols and confidence integration in RegulonDB. Database (Oxford), 2013, bas059, PMID: 23327937

24. Baseggio, N., Davies, W.D. and Davidson, B.E. (1990) Identification of the promoter, operator, and 5’ and 3’ ends of the mRNA of the Escherichia coli K-12 gene aroG. J Bacteriol, 172, 2547–2557, PMID: 1970563

25. Venkatesh, G.R., Kembou Koungni, F.C., Paukner, A., Stratmann, T., Blissenbach, B. and Schnetz, K. (2010) BglJ-RcsB heterodimers relieve repression of the Escherichia coli bgl operon by H-NS. J Bacteriol, 192, 6456–6464, PMID: 20952573

26. Schneiders, T., Barbosa, T.M., McMurry, L.M. and Levy, S.B. (2004) The Escherichia coli transcriptional regulator MarA directly represses transcription of purA and hdeA. J Biol Chem, 279, 9037–9042, PMID: 14701822

27. Myers, K.S., Yan, H., Ong, I.M., Chung, D., Liang, K., Tran, F., Keleş, S., Landick, R. and Kiley, P.J. (2013) Genome-scale analysis of Escherichia coli FNR reveals complex features of transcription factor binding. PLoS Genet, 9, e1003565, PMID: 23818864

28. Burns, P.B., Rohrich, R.J. and Chung, K.C. (2011) The levels of evidence and their role in evidence-based medicine. Plast Reconstr Surg, 128, 305–310, PMID: 21701348

29. Oughtred, R., Rust, J., Chang, C., Breitkreutz, B.J., Stark, C., Willems, A., Boucher, L., Leung, G., Kolas, N., Zhang, F. et al. (2021) The BioGRID database: A comprehensive biomedical resource of curated protein, genetic, and chemical interactions. Protein Sci, 30, 187–200, PMID: 33070389

