## Supplementary material 1 for "A Gold Standard for Transcription Factor Regulatory Interactions in *Escherichia coli* K-12: Architecture of Evidence Types"

The anatomy of the Regulatory Interactions before the additions of evidence derived from the mapping to the HT-TFBSs collection.

This supplementary material file includes the flux diagram for the addition of high throughput (HT)-binding evidence accordingly to the data in the HT-TFBSs collection, as well as additional graphs of the analysis of the RegulonDB version 12.1 regulatory interactions (RIs) set.

Figure S1. Diagram illustrating the process for mapping the set of RIs to the HT-TFBSs collection and the subsequent allocation of HT binding evidence to the set of RIs and the final analyses of their distribution. Self explanatory. All scripts and computational processes built to generate the data and analyses presented in this paper are publicly available and can be found at [https://github.com/PGC-CCG/supplementary-material/tree/master/gold-standard](https://github.com/PGC-CCG/supplementary-material/tree/master/golden-standard)

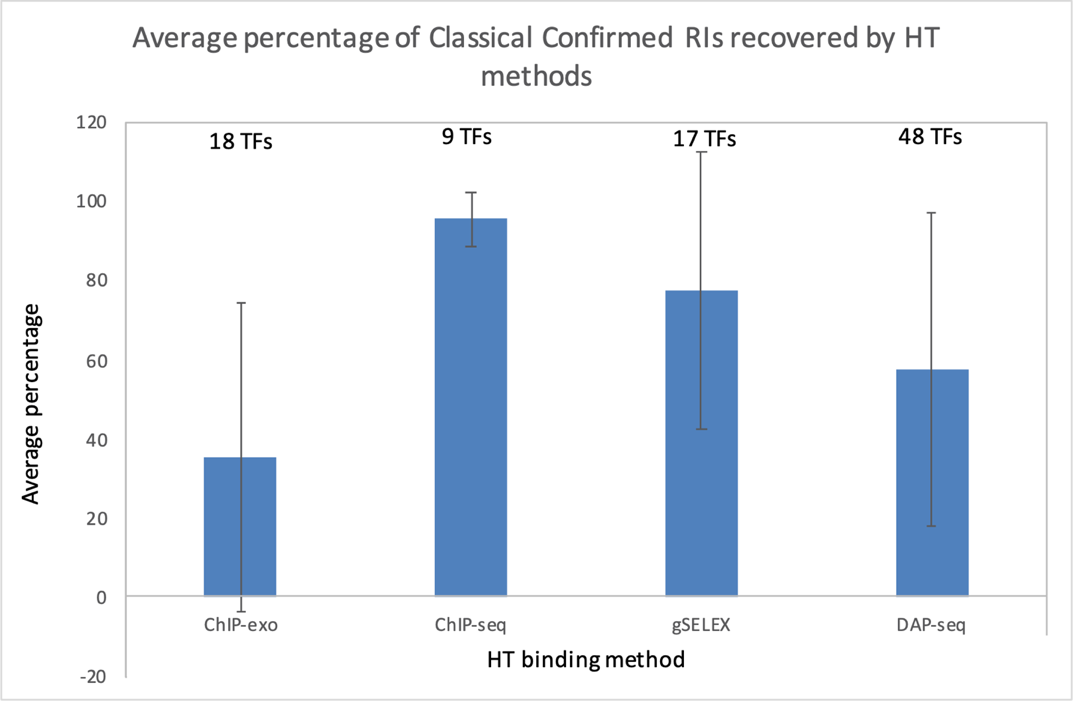

**Figure S2**. Percentage of classical confirmed RIs (RIs that have confidence level “confirmed” which was calculated excluding HT evidence) that were detected by HT binding methodologies. For each methodology the analysis corresponds to a different subset of TFs that meet the requirements of: a) possessing at least one HT dataset and b) have at least one classical RI with confidence level confirmed in RegulonDB 12.1. For each methodology, the fraction of recovered classical RIs was obtained for each TF, and all these were averaged. This figure shows similar results as those shown in Figure 5 in the main paper, where no filtering on confidence level was made.

| **Table S1. Counts of RIs with classical, HT or not experimental evidence and grouped by type.** | | | | |
| --- | --- | --- | --- | --- |
| **RI_type** | **Total** | **Approach** | **# RIs before mapping HT-TFBSs datasets** | **# RI** **s after mapping HT-TFBSs datasets** |
| TF-promoter | 3954 (72%) | Classical | 2426 (61.4%) | 1581 (40.0%) |
|  |  | HT | 93 (2.4%) | 333 (8.4%) |
|  |  | Classical & HT | 248 (6.3%) | 1093 (27.6%) |
|  |  | not experimental | 1187 (30.0%) | 947 (24.0%) |
| TF-TU | 265 (5%) | Classical | 56 (21.1%) | 42 (15.8%) |
|  |  | HT | 156 (58.9%) | 158 (59.6%) |
|  |  | Classical & HT | 20 (7.5%) | 34 (12.8%) |
|  |  | not experimental | 33 (12.5%) | 31 (11.7%) |
| TF-gene | 1247 (23%) | Classical | 19 (1.5%) | 17 (1.4%) |
|  |  | HT | 1107 (88.8%) | 1110 (89.0%) |
|  |  | Classical & HT | 28 (2.2%) | 30 (2.4%) |
|  |  | not experimental | 93 (7.5%) | 90 (7.2%) |

The fourth and fifth columns show the number of RIs of each subgroup before and after to add HT-evidence from HT-TFBSs datasets, respectively. The percent value is relative to the total number of RIs of the corresponding type, indicated in the fourth column.
